# A molecular arm: the molecular bending-unbending mechanism of integrin

**DOI:** 10.1101/2023.04.10.536312

**Authors:** Zhenhai Li

## Abstract

Integrin conformational change is the key to transmitting signals across the cell membrane. The integrin on the cell surface undergoes bending-unbending cycles while sensing and responding to the mechanical environment. Mechanical force triggers the unbending of integrin. However, how an integrin stably extends and how an extended integrin spontaneously bends back are unclear. I performed molecular dynamics simulations on integrin and its subunits to reveal the bending-unbending mechanism of integrin at the atomic level. According to the simulations, the integrin structure works like a human arm. The integrin α subunit serves as the bones, while the β leg serves as the bicep. Thus, the integrin extension results in the stretching of the β leg, and the extended integrin spontaneously bends as a consequence of the contraction of the β leg. This study provides new insights into the mechanism of how the integrin secures in the bent inactivated state and sheds light on the mechanism of how the integrin could achieve the stable extended state.

**Author summary:** As a mechanosensitive protein, the integrin is a molecular machine, which converts the mechano-signal to others, or in the opposite way. The reversible conformational change of integrin is the key to the mechanosensing function. It is straightforward that the integrin would extend while it senses pulling force. However, how integrin bends back after releasing the force is not clear. The ability to bend back guarantees that the integrin can be reused in the bending-unbending cycles. With molecular dynamics simulations, this study shows the integrin works as a molecular arm in the bending-unbending working cycles, which answers how the integrin spontaneously bends back. Meanwhile, based on this study, several hypothesized mechanisms for integrin to be stably extended are proposed.

## Introduction

The integrin is one of the most important transmembrane proteins on the cell surface. It is expressed in multiple different types of cells recognizing glycoprotein ligands in the extracellular matrix or on other cells [1]. Meanwhile, the integrin tail inside the cell links to the cytoskeleton through adaptor proteins. Integrin bridges the extracellular matrix with the cytoskeleton and mediates mechanotransduction [2, 3]. It is involved in cell adhesion, detachment, migration, etc. Cyro-EM images suggest that integrins possess multiple conformations, including the bent, the extended closed, and the extended open conformations[4, 5]. The majority of integrins on the cell surface are in the bent conformation, which is an inactive state in terms of binding to ligands, while very few integrins are activated in the extended open conformation[6, 7]. As a mechanosensor on the cell membrane, it senses mechano-stimuli, and then either directly transmits the signal to the cytoskeleton or converts it to chemical signals via its conformational change, facilitating communications across the cell membrane[1, 8, 9]. Favoring the bent conformation secures that the integrin is ready to sense pulling force, and the ability to reverse to the bent conformation grants integrin the reusing capability. Chen et al. observed the reversible force-regulated integrin conformational change on the cell surface via single-molecule experiments [10]. Surprisingly integrin bent against a pulling force of 20 pN in the experiment. Considering the length change during bending was around ∼13 nm, the integrin had done a work of 360 kJ/mol during bending against the force. It was ambiguous where the energy for bending was from, whether from the energy-storing molecules in cells or the energy stored in integrin.

Molecular dynamics (MD) simulations and steered molecular dynamics (SMD) simulations potentially are proper tools to answer these questions. Chen et al. unraveled a force-induced integrin conformational change pathway via SMD of integrin α_V_β_3_ ectodomains [11]. Puklin-Faucher and Vogel revealed that the binding pocket and βA/hybrid hinge are linked by α1 and α7 helixes with the simulation of head domains of integrin α_V_β_3_. The linkage might be the key to bidirectionality[12]. Nagae et al. showed that Ca^2+^ played an important role in the conformational coupling of the ligand-binding pocket and the rest of the protein[13]. Levin et al. found disulfide bond disruption in the β leg promoted the thiol/disulfide exchange[14]. Su et al. found that the mechanical force can strengthen the interaction between the integrin tail and talin[15]. Lately, researchers have focused on the full-length integrin embedded in the cell membrane. Owing to the computational limitation, all-atom simulation on the full-length integrin is time-consuming. Therefore, coarse-grained MD simulations were performed to explore various conformations of the full-length integrin embedded in the cell membrane[16]. However, it is still unclear why the integrin favors the bent inactivated conformation and how integrin spontaneously bends back.

Herein all-atom MD and SMD simulations on integrin α_V_β_3_ ectodomain were carried out. In the SMD simulations, pulling force was applied to bent integrin α_V_β_3_ to induce the extension. Then the MD simulations were performed, in which the force was released to examine the spontaneous bending of the extended integrin. In the simulation, integrin worked like a molecular arm. The leg in the β subunit served as the bicep in the arm. The β leg was stretched during the force-induced extension. After the force was released, the contraction of the β leg triggered the spontaneous bending of integrin. Besides β leg stretching, the integrin head-to-leg interaction was disrupted with integrin extension. Thus, the extended integrin stored energy in the β leg and the head-to-leg interface. Based on the theoretical calculation and umbrella sampling simulation [17], the stored energy in the extended integrin is estimated over 500 kJ/mol, which covers the work required for integrin to bend back against a force even over 30 pN.

## Results

### The summary of simulations

The MD and SMD simulations were performed on three different constructs: integrin ectodomains, α subunit alone, and β subunit alone. The simulation model for the whole integrin and the subunit alone are all from the crystal structure of α_V_β_3_ (Protein Data Bank code: 3ije). The whole integrin or subunits alone were positioned in a water box with the size of 16×17×30 nm^3^. Na^+^ and Cl^−^ were added to achieve the 150 mM salt concentration and neutralize the system. In total, the integrin simulation system includes 542,742 atoms (Fig. 1A). As the α and β subunits structures were obtained from the integrin complex, 20-ns initial MD simulations on these two constructs were applied to obtain the relaxation structures for the subunits. Then the whole integrin and the subunits alone were pulled with 4∼5 independent SMD simulations for 15 ns, respectively. In the SMD simulation, the Cα atom of VAL681 in the β leg (or VAL951 of α leg for the α subunit alone simulation) was constrained, and a group of atoms close to the integrin-RGD binding interface was pulled with a harmonic spring at 50 kJ/(mol·nm^2^) (see details in Methods). The spring was moving at a constant velocity of 1 nm/ns. After the pulling, 50-ns MD simulations were applied to whole integrin and subunits alone, in which the force was removed allowing the bending of integrin.

**Fig. 1.**
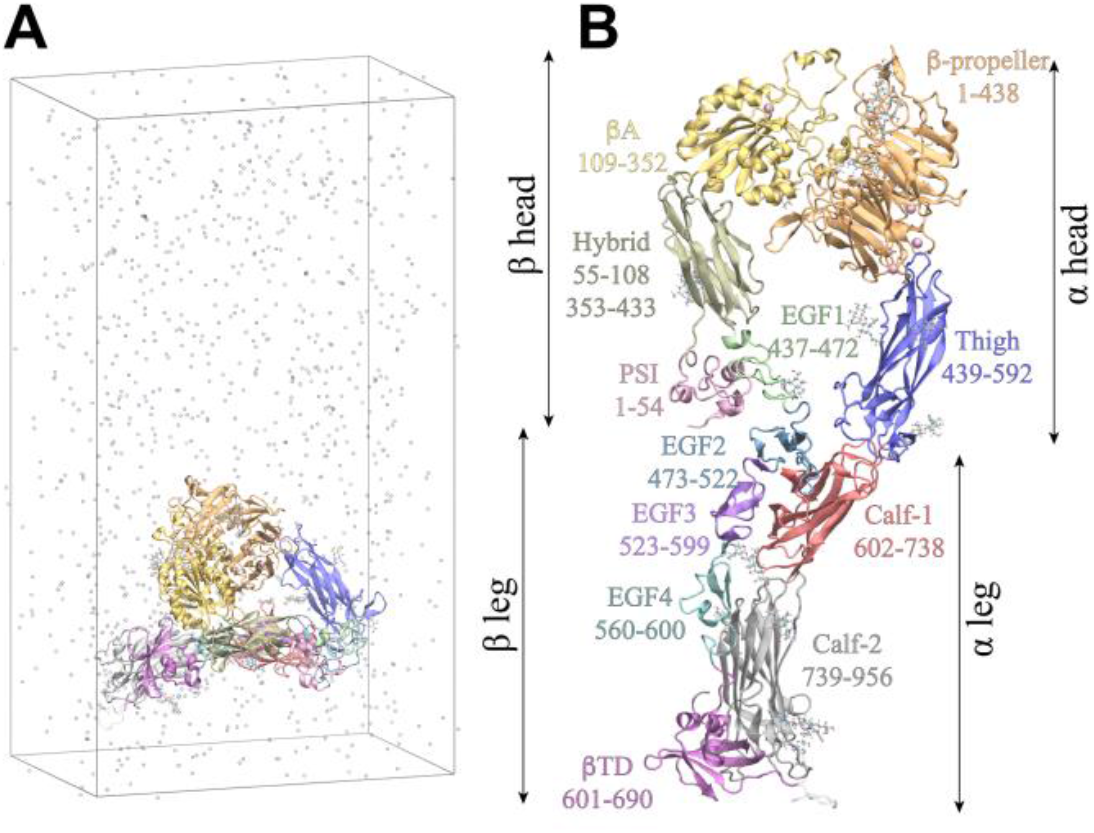
The simulation system and the integrin domain definition. A. The integrin in the simulation box with solvent (only Na^+^, Cl^−^, and Ca^2+^ are shown and the water molecules are not shown for the clarity). B. The amino acid range of each domain in two subunits.

Since the pulling was at a high loading rate, the conformation obtained by SMD might not be at equilibrium. To avoid the integrin promptly bent back due to the tension created by the fast pulling, four sets of additional simulation runs were performed. In the additional simulations, equilibrium simulations with length constraints were carried out for different time periods, and then another 200-ns MD simulation without any constraints was applied to allow the integrin bending. To examine the energy landscape of integrin during extension, umbrella sampling (US) simulations were carried out on the integrin, in which, 47 configurations generated along a SMD simulation were used. The total simulations time, including SMD, MD, and US simulations, is over 4 μs.

### Integrin unbending and bending

A modeled extended integrin is shown in Fig. 1B to clarify the domain definition of integrin. Integrin α subunit includes a β-propeller, a Thigh, a Calf-1, and a Calf-2 domains. The first two domains are in the α head and the last two are in the α leg. The integrin β subunit includes a βA, a hybrid, a PSI, four EGF (including EGF1, EGF2, EGF3, and EGF4), and a βTD domains. The first 4 domains including βA, hybrid, PSI, and EGF1 are in the β head and the rest domains are in the β leg (Fig. 1B).

In the simulation, the distance between the center of mass of the pulling group and the constrained Cα was defined as the integrin length. A slight increase in the length of the α subunit and a slight decrease in the length of the β subunit were observed in the initial relaxation. With the force applied to the integrin head, the whole integrin, the α subunit, and the β subunit all extended and the total length increased by ∼13 nm (from ∼5 to ∼18 nm), ∼18 nm (from ∼7 to ∼25 nm), and ∼14 nm (from ∼4 to ∼18 nm), respectively (Fig. 2A, B). Besides the unbending of the head and leg, a clear extension in the β leg EGF domain was observed in the β subunit simulation (right panel in Fig. 2A).

**Fig. 2.**
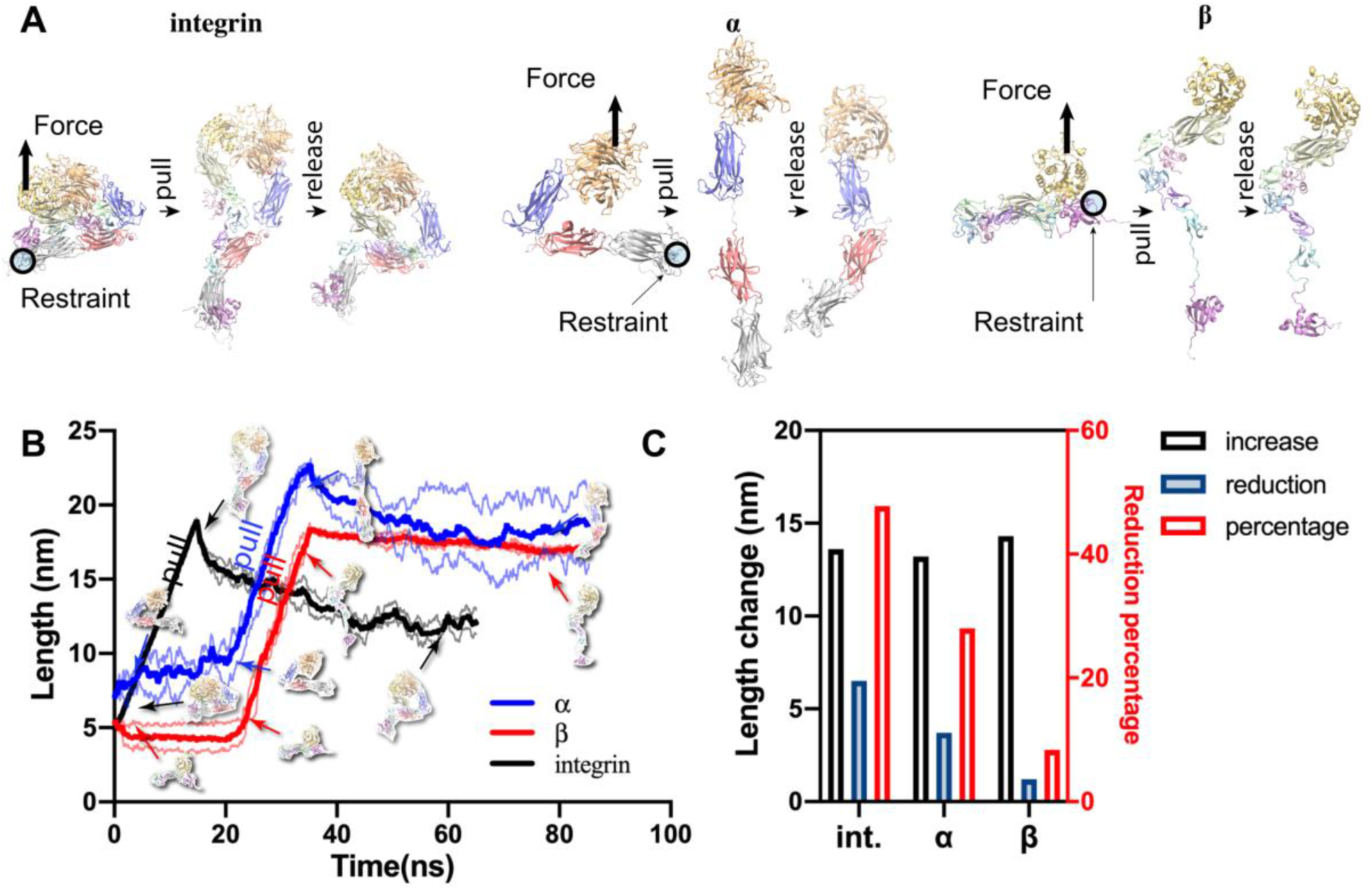
Pulling and bending simulation of full integrin and two subunits. A. snapshots of pulling and bending simulations of integrin (left), α subunit (middle), and β subunit (right). B. the length of integrin or α/β subunit alone during pulling and bending simulation. The insets are the snapshots of integrin, α, or β subunit structure during the simulation. The thick lines are the average length over 6 independent simulations, the thin lines are the upper and lower range of these simulations. C. the length increase from the relaxation to full-extended state and the length reduction from the fully-extended to the force-released states. The reduction percentage shows the ratio of reduction to increase.

Afterward, the force was released, and the lengths of all three constructs decreased to different extents (Fig. 2B, C). In Fig. 2C, the length increase referred to the length change from the relaxation to the fully extended state. The length reduction referred to the length decrease from the fully extended state to the final state with force released. The length extension was eventually reduced respectively by 50%, 30%, and 20% for whole integrin, α subunit, and β subunit after the force was released (Fig. 2C and Supplementary Video S1-S3). The length reductions for the three constructs were different. Although the head and leg region of the β subunit has the potential to form extensive interactions[18], the interaction interface in the head and leg were far from each other in the extended β subunit. Therefore, the interaction failed to form and the β subunit stay extended. The length reduction of the β subunit alone is mainly from the shrinking of the EGF domains in the β leg (Supplementary Video S2). Intriguingly the crystal structure of the β subunit alone is also in an intermediate or the extended conformation [19], which agrees well with the simulation results. The length reductions of the α subunit alone and whole integrin were from the bending of the head and leg region. However, the α subunit bent to the opposite side (middle panel in Fig. 2A). Moreover, a salt-bridge of ASP457−LYS688, an interaction between ASP599, GLU636, and Ca^2+^ at the genu of α subunit formed, which could lock the α subunit in the opposite bent conformation (Supplementary Fig. S3 and Supplementary Video S3).

### The synchronized length changing of integrin and its β leg

Intriguingly, the integrin complex could bend back after the force is released although neither of the subunits is able to do so. In order to inspect the mechanism for integrin bending, the deformations of head and leg regions of α and β subunits in both the SMD and the MD simulations on the whole integrin were examined (Fig. 3A). The distance between the Cα of ASP219 and ILE592 of the α subunit, CYS602 and THR952 of the α subunit, TYR122 and GLU472 of the β subunit, and CYS473 and GLU683 of the β subunit were respectively defined as the α head, α leg, β head, and β leg length. The integrin length increased from ∼5 nm to ∼18 nm during pulling, meanwhile, the β leg length increased by ∼2 nm (blue curve in Fig. 3A and Supplementary Fig. S1). After the force was released the β leg was contracted to the relaxation length, which was followed by the bending of integrin (Fig. 3A). Moreover, the head region including the α and β subunits had limited changes in length. The α leg was even shortened owing to the bending of the Calf-1 and Calf-2 domains, suggesting the α leg was under compressive force (Fig. 3A).

**Fig. 3.**
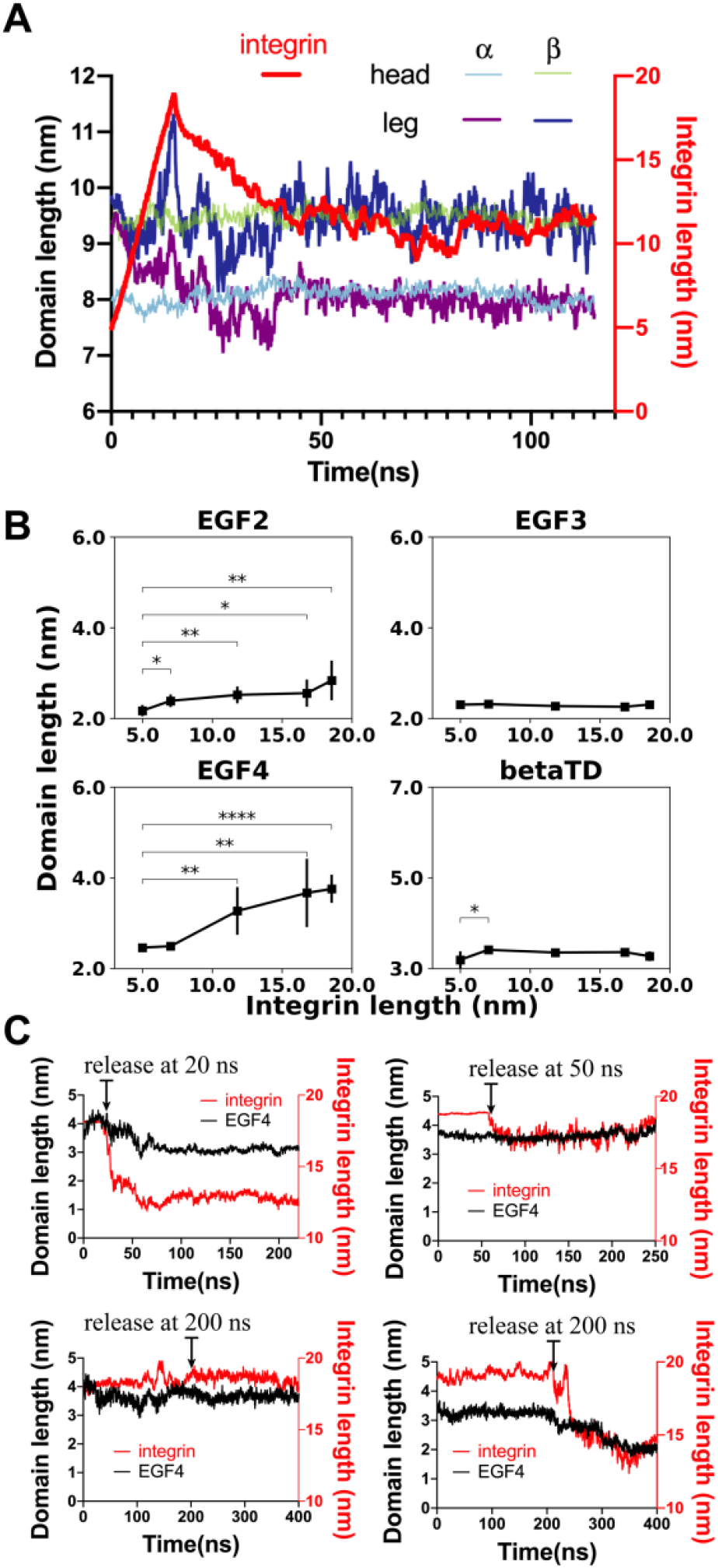
Deformation of the integrin during unbending. A. the integrin head/leg region of α/β subunit length changing with integrin total length changing over time. B. the length of domains in the β leg at different integrin lengths. C. Prolonged simulations, the integrin after being pulled in the pulling simulation were clamped at the extension state and then released at a specific time point as indicated in each panel.

To quantify the deformation of each domain, the simulation trajectories were aligned to each domain and then the root mean square deviation (RMSD) of each domain over time was calculated. According to the RMSD, the greatest deformation occurred in the EGF4 domain in the β leg (Supplementary Fig. S2), EGF1, EGF2, and EGF3 had moderate RMSD increasing during pulling. The RMSDs of the rest of the domains remained at the level of ∼0.3 nm, indicating limited deformation occurred in these domains during the pulling. The RMSD of each domain suggested the α subunit and β head region are relatively rigid, while the β leg is relatively flexible (Supplementary Fig. S2).

As the β leg is most flexible, the length changes of domains in the β leg were further studied. Five integrins at different lengths (including bent, fully extended, and three partially extended as in Fig. 3B) were obtained from the pulling simulation. Then, an 8-ns MD simulation was performed for each integrin structure. In each simulation, the integrin length was clamped by restraining the pulling group and the Cα of V681 in the β leg. The integrin total length and the length of each domain in the β leg were calculated. The simulation with restraints at each certain length was repeated 5 times independently, and the average domain lengths and integrin length were then calculated over 5 simulations. The EGF4 length significantly increased with integrin length, while the EGF2 length increased slightly (Fig. 3B).

To reduce the possible artifact introduced by the high-speeding pulling, the integrin was clamped after full extension for 20 ns, 50 ns, and 200 ns (as indicated in Fig. 3C) by applying restraint at the pulling group and the Cα of V681 in β leg to find equilibrated conformation, afterwards 200-ns MD simulations were performed with the restraints released (Fig. 3C). In the simulation with 50-ns clamping and one of the simulations with 200-ns clamping, spontaneous bending was not observed in the following 200-ns simulation without any restraints. In the other two simulations, spontaneous bending occurred. Intriguingly, in the two simulations without integrin bending, the EGF4 domain length failed to shrink either, while in the other two simulations, the spontaneous bending occurred with the length contraction of the EGF4 domain.

### An important interaction between Thigh and EGF2 domains

Another key observation was a stable interaction, which drew less attention previously, existed between the EGF2 in the β leg and the Thigh domain in the α head. The EGF2–Thigh interaction plays an important role in the integrin conformational change. It has been reported there were extensive interactions including the hydrogen bond (H-bond) and hydrophobic interaction between the integrin β head and β leg [18, 20]. These interactions were disrupted during the pulling (Fig. 4A, B). However, at the EGF2–Thigh interface, there are four basic amino acids (K501, K503, R509, R549) and four acidic amino acids (D512, E500, E475, E476) on the Thigh and EGF2 domains, respectively. These amino acids formed ∼2 salt bridges on average in the free dynamics simulations. Surprisingly, unlike the interactions between the β head and leg, which dissociate upon pulling, these two salt bridges survived throughout the pulling (Fig. 4C, D). It is worth noting that the EGF2–Thigh interaction is very close to but off the hinge by 1∼2 nm.

**Fig. 4.**
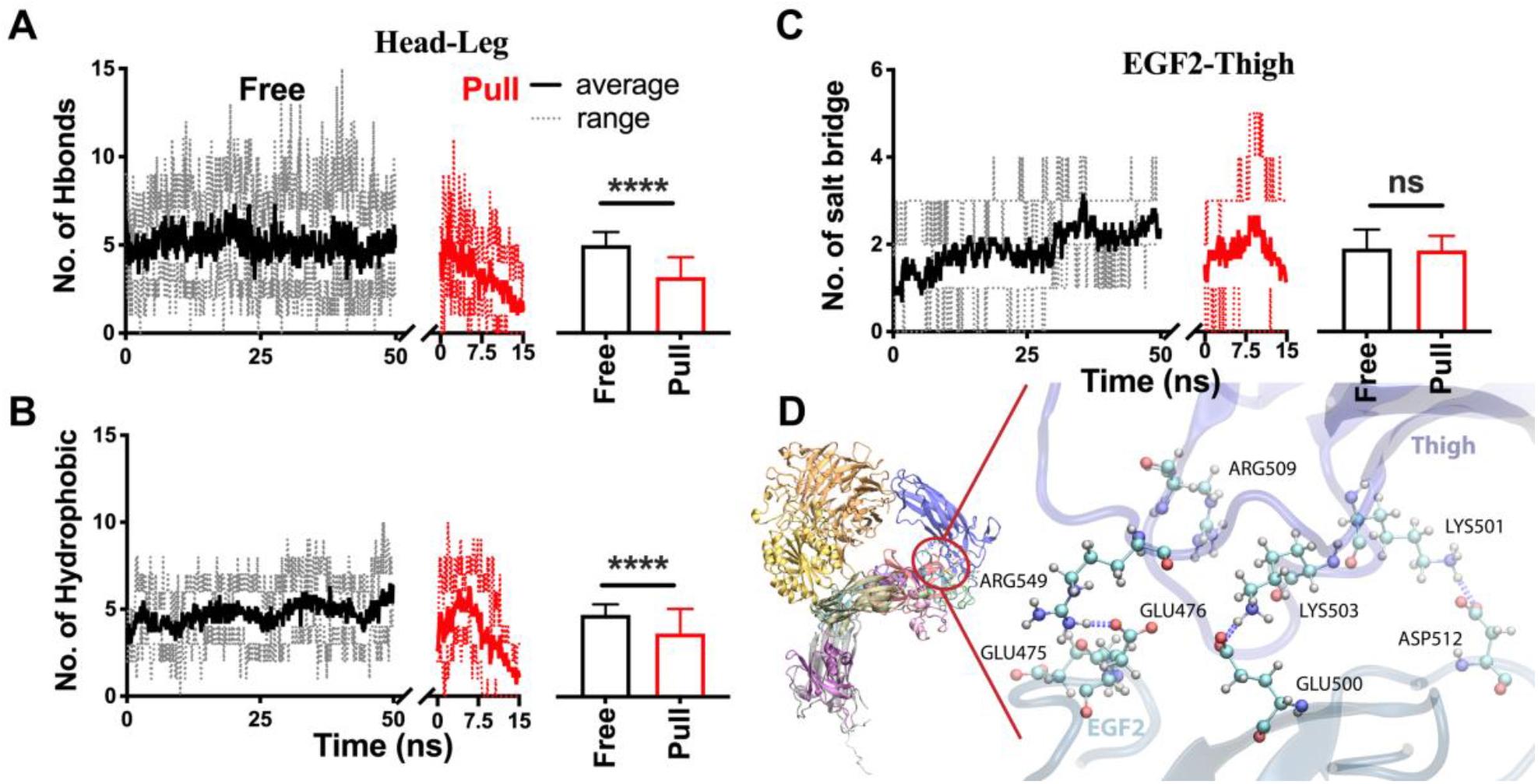
The interaction between Thigh and EGF2 domains. A. the numbers of H-bond and hydrophobic interaction between integrin head and leg change during free dynamics and pulling simulations. B. the primary salt bridge between EGF2 and thigh domain. C. the number of the salt bridge in the free dynamics and pulling simulations.

### The arm-like working mechanism

These two observations, the β leg and integrin synchronized length changing and the off-hinge interaction of EGF2–Thigh, reminded me of the human arm working mechanism (Fig. 5). One can use a metaphor to easily understand the integrin bending mechanism. In the metaphor, the integrin bending can be simply described as an analog to the arm. The forearm is analogous to the integrin head; the upper arm is analogous to the integrin leg; the bones in the arm are analogous to the hard α subunit; the bicep is analogous to the integrin β leg. The bicep is attached to the tuberosity in the forearm close to but off the elbow joint, which is similar to the off-hinge interaction between the β leg and the Thigh domain. When the bicep crunches, the lower arm is brought back and induces the bending of the arm. Similarly, when the β leg is contracted, the integrin bends back.

**Fig. 5.**
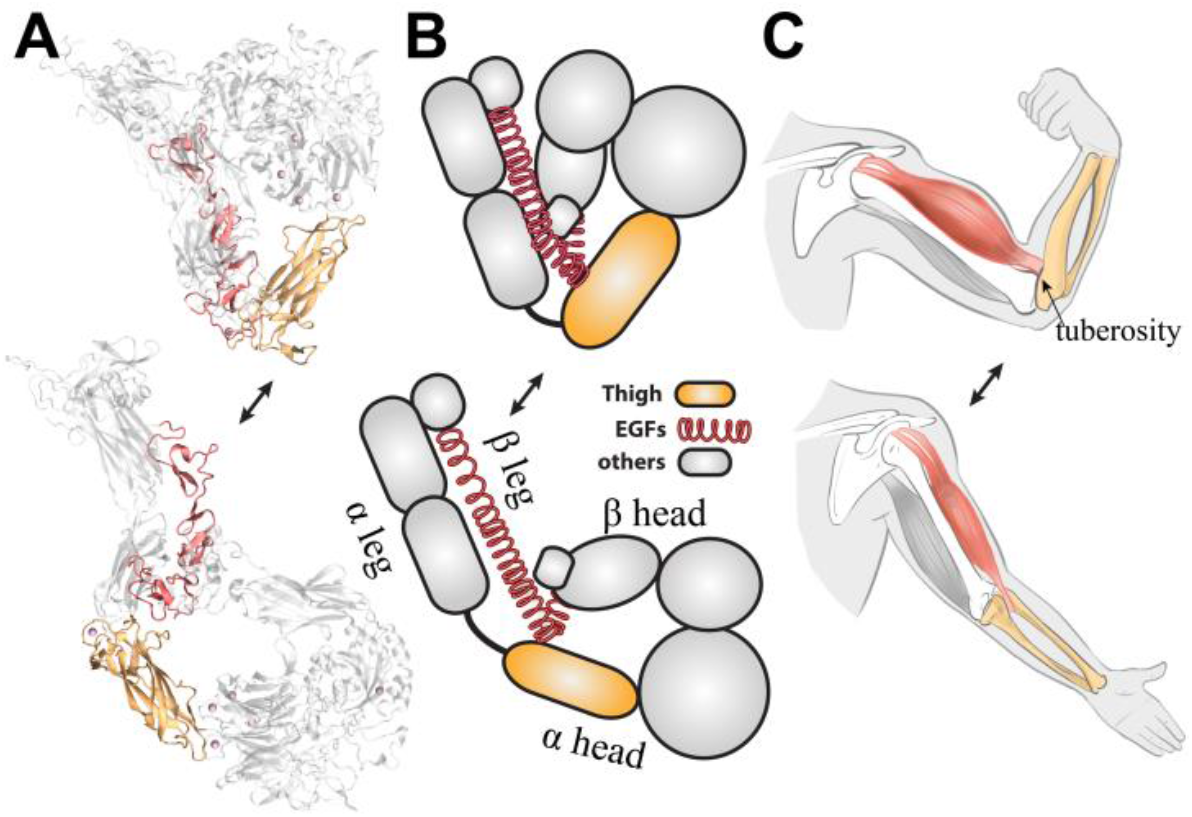
integrin resembles a human arm. A. the simulation snapshot of integrin pulling. B. a simplified model of integrin conformation change. C. human arm bending-unbending mechanism.

A prerequisite of the arm bending is the bicep attaching to the tuberosity of the forearm. If the muscle is peeled off from the bones, the contraction of the bicep would no longer induce bending of either muscles or bones. Similarly, if the α and β subunits are separated, neither of them can bend back from the extended conformation as observed in the SMD and MD simulations of the α and β subunits alone (Fig. 2 and Supplementary Video S2 and S3).

### The energy landscape of integrin

If the integrin conformational change does follow the mechanism similar to the arm, considering the α head and leg as rigid rods, and the β leg as an elastic string, the integrin structure can be simplified as a trusswork (Fig. 6A). According to the geometric relationship, the β leg and integrin total length would follow:

**Fig. 6.**
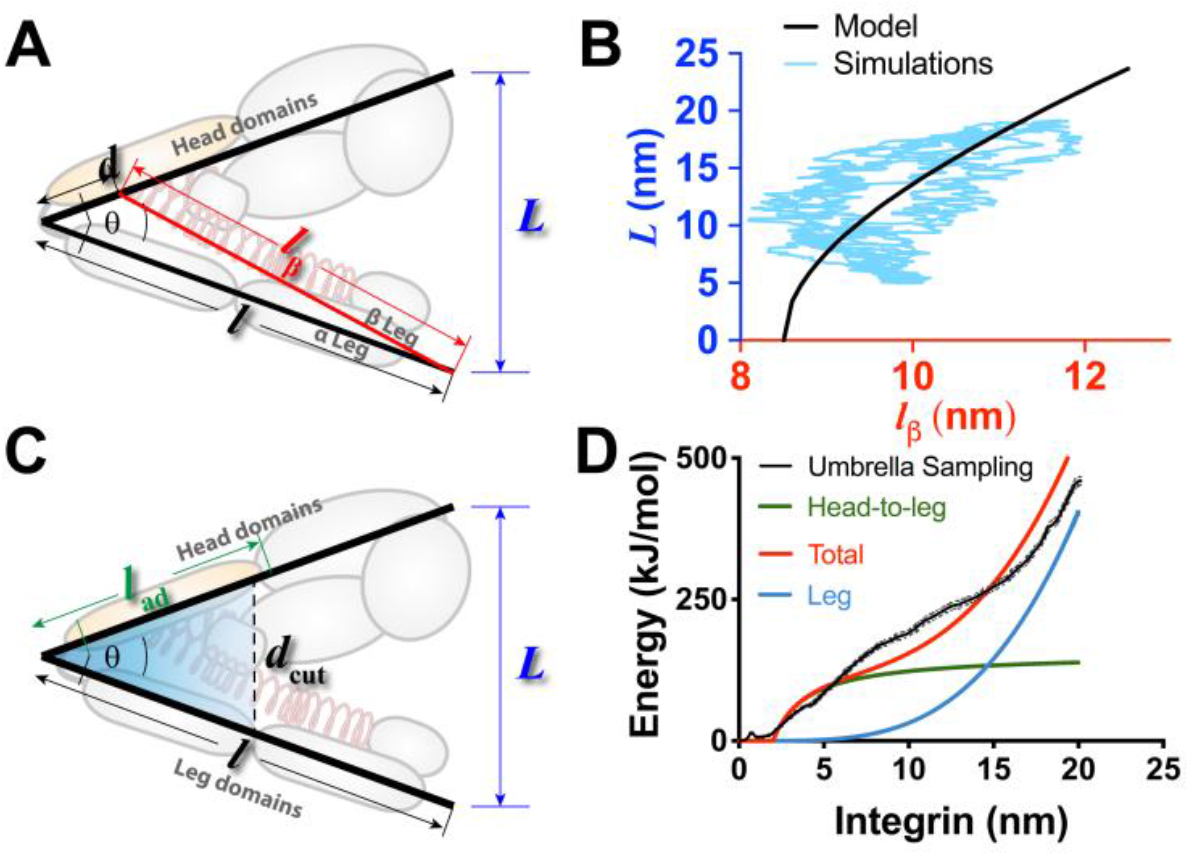
The energy landscape estimation. A. trusswork of integrin. B. the correlation of integrin length and β leg length. C. the model to estimate the integrin head-to-leg contact area. D. integrin pulling energy landscape, the standard deviation of the energy obtained by Umbrella Sampling simulation were indicated by the dashed line.

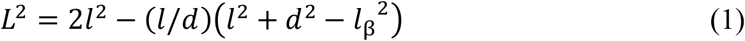

in which the α head and leg lengths are equal and denoted by *l, l*_β_stands for the β leg length, and *d* stands for the distance from the off-hinge interaction site to the hinge (Fig. 6A). All these characteristic lengths can be measured in the crystal structure, where *l*≈ 10 nm, *d* ≈ 1.5 nm. *l*_β_varies during the pulling. As *l*_β_changes, the total integrin length *L* changes (Fig. 6B). This geometric relation agrees well with the observations in the steered MD simulations (Fig. 6B).

From the energy perspective, new surfaces are created with the head-to-leg interaction disruption, and then the β leg is stretched during integrin extension. Thus, the energy elevated is primarily stored as the surface energy in the head-to-leg interface and the elastic energy in the stretched β leg. During pulling, ∼5 H-bonds and ∼5 hydrophobic interactions are broken between the head and leg pieces with the integrin extension (Fig. 4A, B). The energy of a single H-bond is around ∼7.3 kJ/mol [21]. The energy of the hydrophobic interface is around 36 kJ/(mol· nm^2^) [22]. The hydrophobic surface change during the unbending of integrin is obtained by calculating the solvent accessible surface area (SASA) of hydrophobic amino acids in the MD simulation (Supplementary Fig. S4). It increases by 6.5 nm^2^ during the pulling. Adding up the H-bond and hydrophobic energy, the total surface energy increase is 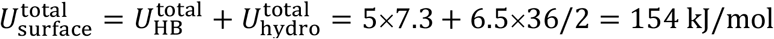. Considering the head and leg interfaces maintain interaction if their distance was within a cutoff distance, *d*_cut_, the interacting region can be calculated as *l*ad = min [*d*_cut_ · (*l*/*L*), *l*] (Fig. 6C). The H-bond cutoff is 0.3 nm in general. However, integrin is flexible, especially the β subunit which makes extensive contact between the head and leg. Thus, it is reasonable to loosen the cutoff. I choose 2 nm as *d*_cut_ for the disruption of the head-to-leg interaction in the model. Assuming the interaction between the head and leg is equally distributed along the interface with an energy density, 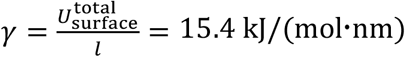, the elevated head-to-leg adhesion energy with integrin extension can be calculated by *u*_surface_ = *γ*(*l* − *l*_ad_).

The energy stored in the integrin leg can be simply described by Hooke’s law as: *u*_leg_ = 1/2 · *k*(*l*_β_− *l*_β_0)2, where *k* is the spring constant of β leg, and *l*β0 = 8.5 nm is the β leg at relaxation while integrin is at the bent conformation. The spring constant of the β leg can be calculated based on the directional variance of the protein in the MD simulation trajectories [23]. The estimated spring constant is ∼90 kJ/(mol·nm^2^).

The total energy change of integrin during the extension of integrin can be calculated by the following master equation:

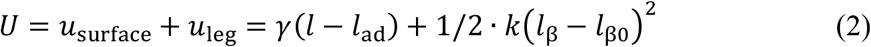

By replacing *l*_ad_ and *l*_β_ with *L*, the relation between integrin energy change and total length is established. Using the parameters above, the energy change vs. total length was calculated (Fig 6D). The free energy does not change with the length change at the first 2 nm. This is for two reasons: 1. at the beginning of the pulling, especially when the integrin length *L* is lower than 2 nm, the greatest head to leg distance is within the cutoff distance, and the interaction between head and leg has not been disrupted yet and the surface energy keeps unchanged; 2. the β leg length increases very slowly at the beginning of integrin extension (Fig. 6B), results in slow increasing of elastic energy in the β leg. Afterward, the energy increases rapidly, which is caused by the fast disruption of the head-to-leg interaction. After the integrin length is over 5 nm, the free energy increasing slows down, until the integrin length reaches ∼15 nm the free energy fast increases again owing to the fast stretching of β leg. Therefore, the energy landscape presents a slow–fast–slow–fast manner.

We further did the umbrella sampling (US) simulation [17] to calculate the free energy change with the integrin extension. 47 configurations along the a SMD simulation were extracted. The adjacent configurations had a ∼0.25 nm integrin length increment. Then the integrins at different lengths were held at the corresponding lengths for 20 ns by applying a harmonica potential to the pulling group. Using Weighted Histogram Analysis Method (WHAM)[24], the free energy landscape was obtained (Fig. 6D). The US-simulated energy increases with the length extension, and the increasing experiences four stages: slow–fast–slow–fast. This free energy landscape agreed well with the model predicted results (Fig. 6D).

## Discussion

The human arm model suggests the contraction of the β leg initiates the spontaneous bending of integrin. In addition, based on the human arm model and previous MD simulation findings, one can propose several possible mechanisms for integrin to stably extend. 1. In a previous MD simulation[11], Chen, et al. showed a metal ion at α genu can interact with the amino acid D457 in α head in the extended conformation, forming a metal ion coordination (Fig. 7A). Bending of integrin will induce high tension force in the metal ion coordination, which can counter the contraction of the β leg and stabilize the extended conformation.

**Fig. 7.**
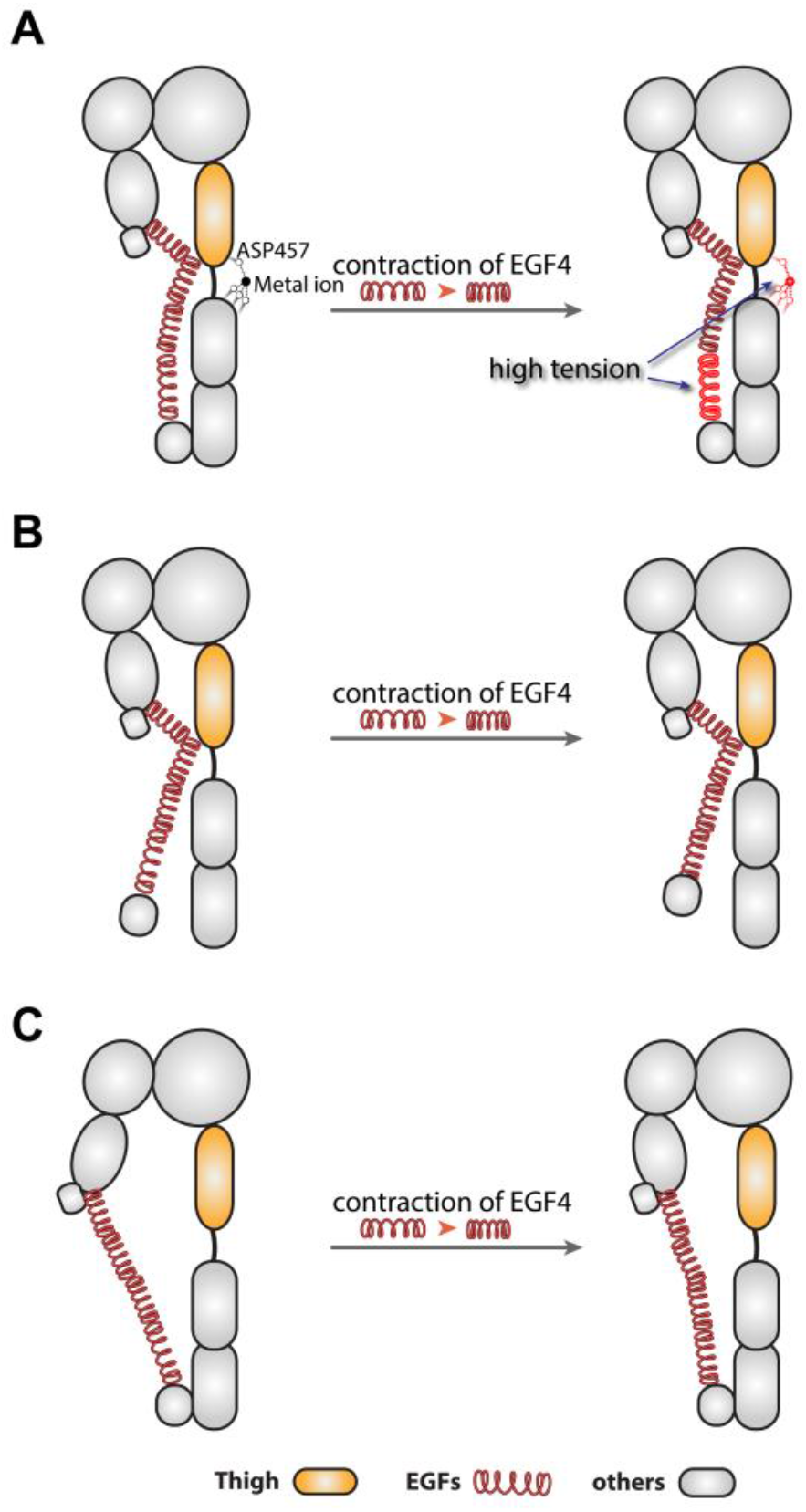
The possible mechanism for integrin extension. A. the metal ion coordination at the α genu. B. splitting the α and β legs. C. hybrid domain swinging out and breaking the EGF2– Thigh interaction.

2. The splitting of legs possibly promotes the extended conformation (Fig. 7B). If the interaction between the α and β legs is disrupted, the contraction of the β leg can no longer drag the integrin head region to bend back, instead, the β leg will be freely shortened. The current view believes the adaptor protein, talin, binds to integrin β leg cytoplasmic tail and pulls the β tail away from the α subunits. The disruption of the α and β interaction induces the extension and activation of integrin[25, 26]. The human arm model provides a possible explanation of how the disruption of α and β facilitates the integrin conformational change and activation.
3. The disruption of EGF2–Thigh interaction may promote the integrin extended conformation (Fig. 7C). If the interaction between the EGF2 and Thigh domains is disrupted, the contraction of the β leg can no longer drag the integrin head region back either. The shortening of the β leg may only rearrange the orientation of each domain in the β subunit but cannot induce bending of integrin (Fig. 7C). Smagghe et al. introduced mutations to the EGF2 domain of integrin, in which multiple amino acids from CYS473 to CYS486 were replaced. The authors concluded the β knee served as an entropic spring, which favored the bent conformation. The shortening of the loop between CYS473 and CYS486 altered the balance of the entropic spring and induced the integrin extension[27]. However, a recent study demonstrated that the integrin β subunit alone exhibited an extended conformation[19], suggesting the entropic spring may not exist. It’s worth noting that in Smagghe’s study GLU475 and GLU476 were mutated, which may weaken the interaction between EGF2 and Thigh (Fig. 4D). As the human arm model suggested, the weakening of EGF2–Thigh interaction may also facilitate the extension of integrin. Thus, the bicep model provides another possible explanation for Smagghe’s results. In addition, the hybrid domain swing-out is involved in integrin activation[4, 28]. Crystallographic and computational studies have unraveled that the subtle structure change in the βA domain occurs with the swing-out of the hybrid domain, which potentially enhances the binding affinity to the ligands[29, 30]. According to the human arm model, the hybrid domain swing-out also disrupts the EGF2–Thigh interaction, which could also stabilize the extended conformation.

Based on the bicep model, the contraction of the integrin β leg, especially the EGF4 is the driving force of the integrin bending. Therefore, the flexibility of the β leg is critical to the bending-unbending conformational change. There are three EGF (EGF2 to EGF4) domains in the leg, each EGF domain contained four possible disulfide bonds. The disruption of the disulfide bond in EGF domains can increase its flexibility. As a result, the extended conformation would be enhanced. Several studies have shown the disulfide bond can regulate the integrin structure and function[31-33]. A previous study suggested that the mutation of cysteines in EGF domains promoted the integrin activation, but the mutations of cysteines in other domains of the β subunit did not. However, the molecular mechanism is unclear[34]. According to the human arm model, the mutation of cysteine in EGF domains disrupt the disulfide bond in EGF domains, which increases the flexibility of the β leg, promoting the extension of integrin and facilitating its activation.

Integrin on cell surface experiences bending-unbending cycles. The integrins at the leading edge of the cell bind to the matrix through interaction with their ligands and link to the cytoskeleton through the interaction between the β tail and adaptor proteins. The force generated by the cytoskeleton pulls the integrin at the leading edge, potentially activates integrins, and meanwhile drags the cell to move forward. As a result, the integrins translocate to the proximal end which is close to the center of the cell, and eventually dissociate there. In the cycle, the inactive integrin senses force and gets activated, and eventually gets deactivated when force is released. This MD simulation and the theoretical model unravel the molecular mechanism to explain why an integrin favors the bent inactive state, which inhibits the unnecessary activation of integrin and gets integrin ready for mechanosensing, and illustrates how an activated integrin spontaneously bends back with the aid of the high elastic energy stored in the integrin β leg and the surface energy, and complete a cycle of the conformational change.

## Methods

### System setup

The integrin α_V_β_3_ (PDB code 3IJE) was used as the initial structure for the simulation. The α and β subunits alone were obtained by removing the other subunit from integrin α_V_β_3_ (3IJE). The full integrin and the subunit alone were placed in a water box with a size of 16×17×30 nm^3^ (Fig. 1A). TIP3P water and 150 mM NaCl was added to the box to mimic the physiological environment (Fig. 1A). The simulation system included 542,742 atoms. The CHARMM22 force field[35, 36] is used to describe the protein. The simulation is carried out with the GROMACS package [37]. An energy minimization with the combined steepest descent and conjugate gradient methods was first applied to the system. Afterward, the system was gradually heated to 310 K within 1 ns and held in NVT for 1 ns and then NPT ensemble for another 1 ns with the protein constrained. The temperature and pressure are controlled by V-rescale [38] and Parrinello–Rahman algorithms [39, 40], respectively. Afterward, the MD and SMD simulations were applied to integrin and its subunits. In the SMD simulation, a group of atoms at the ligand interaction interface were pulled with a spring of 50 kJ/mol/nm at the speed of 1 nm/ns for 15 ns. In the restrained simulation and the US simulation, the pulling group atoms were restrained with the same spring at a moving speed of 0. The pulling group included the heavy atoms in βA residues 113-117, 151-156, 190-197, 244-250, 306-310, 329-332 in β subunit and β propeller residues 22-26, 97-101, 160-164, 225-229, 279-283, 343-347, 407-411 in α subunit. In the simulation on the whole integrin, the pulling group atoms in both subunits were pulled. In the subunit alone simulation, the pulling group atoms in the respective subunit were pulled. For the simulation on the whole integrin and β subunit, the Cα of VAL681 of the β subunit was constrained. For simulation on the α subunit, the Cα of VAL951 of the α subunit is constrained.

### Umbrella sampling simulations

From one SMD simulation of integrin, 47 configurations along the pulling were generated. The adjacent configurations were selected with a length spacing window of 0.25 nm. Then 47 MD simulations were applied to these integrins with the protein being held at their current lengths by applying a harmonic potential with a spring constant of 50 kJ/mol/nm to the pulling group for 20 ns. The energy landscape of integrin along the extension was analyzed by using Weighted Histogram Analysis Method (WHAM). The statistical error was analyzed by Bayesian bootstrap analysis with 10 bootstraps.

