## Supplementary Fig S1, S2, S3, S4 for "A molecular arm: the molecular bending-unbending mechanism of integrin"

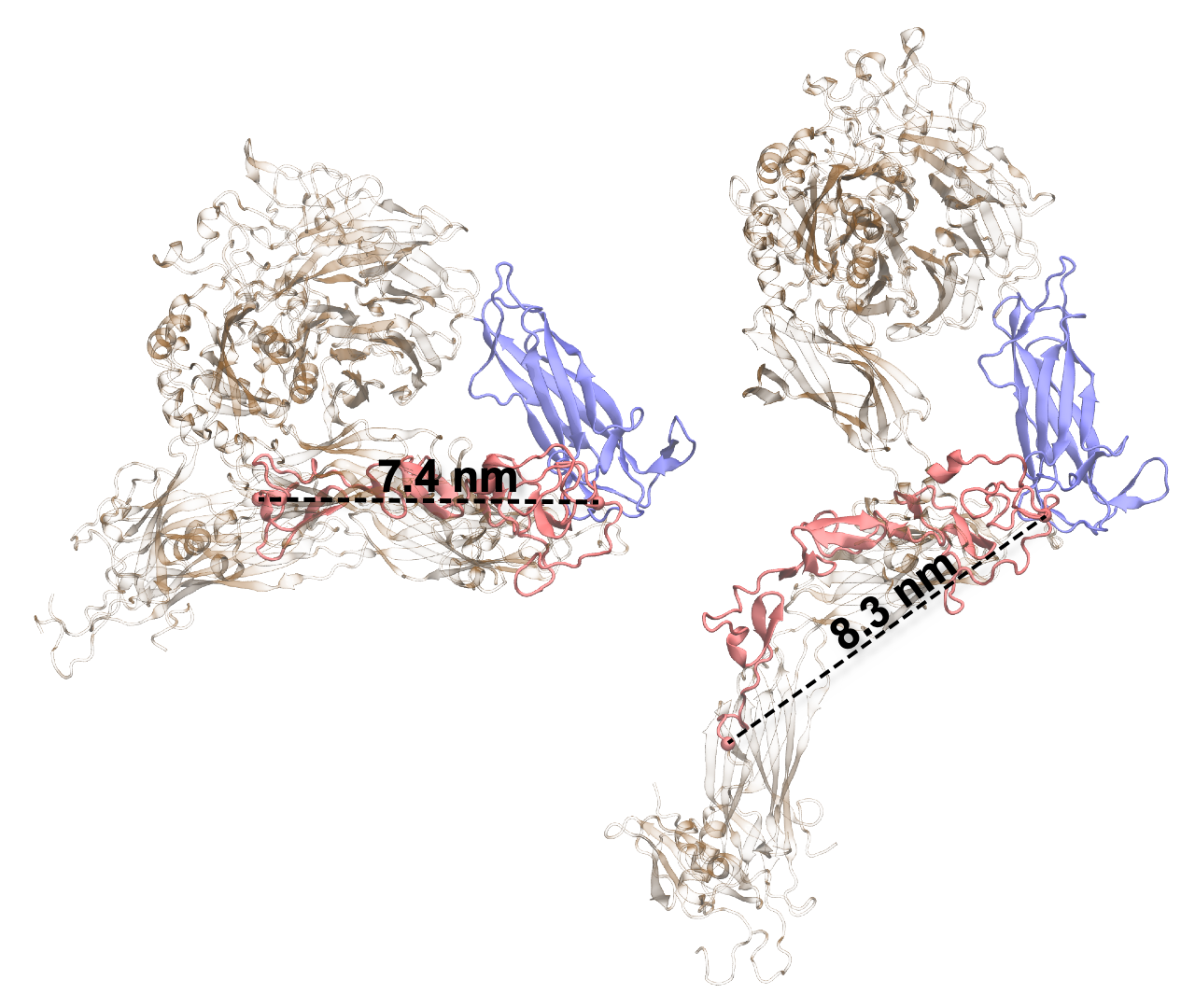

Supplementary Fig. 1. The integrin β leg length increased with the integrin extension.

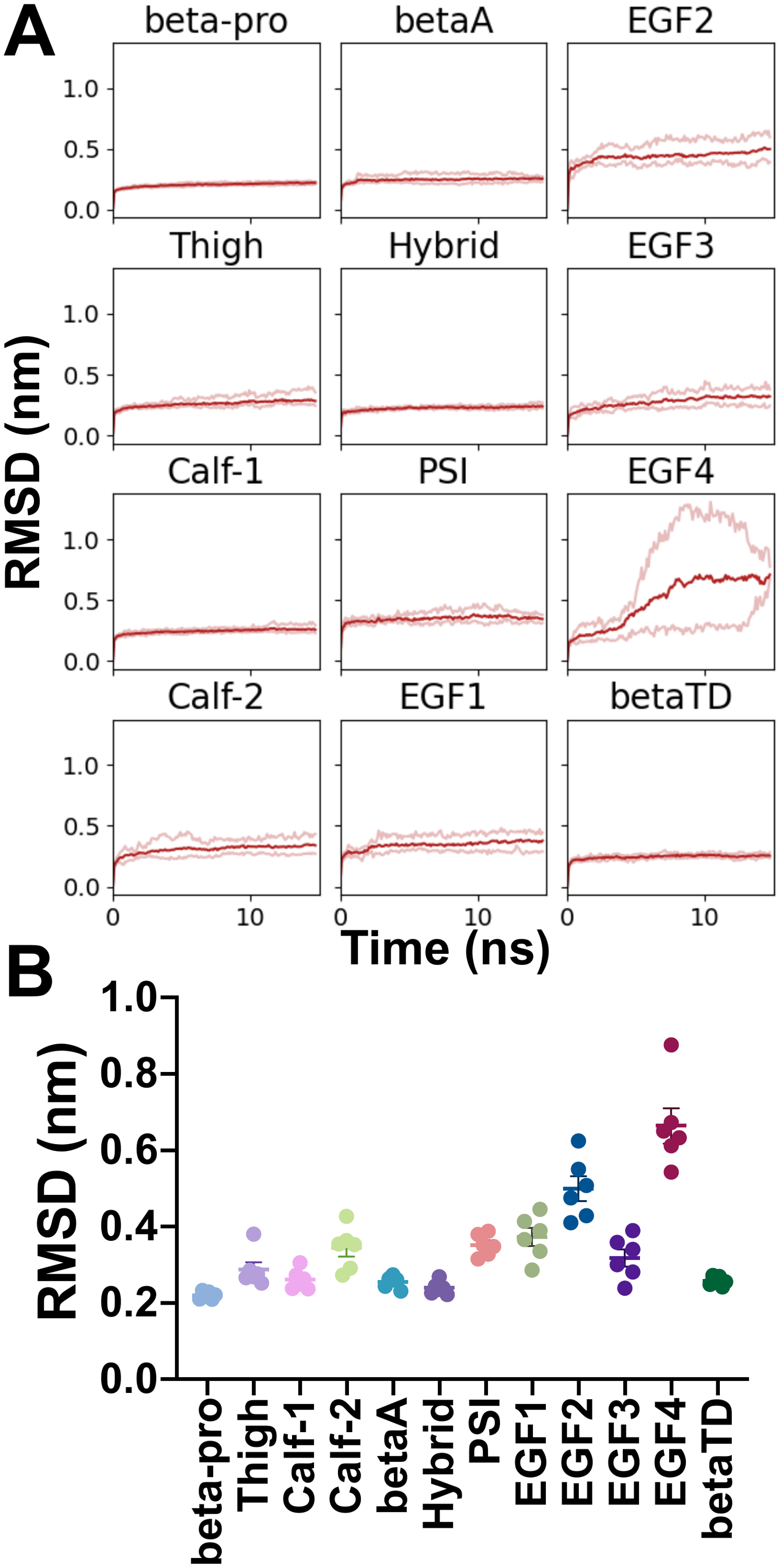

Supplementary Fig. 2. (a) The RMSD of each integrin domain in the 6 independent pulling simulations. (b) the average RMSD in last 10 ns of the 6 independent pulling simulations.

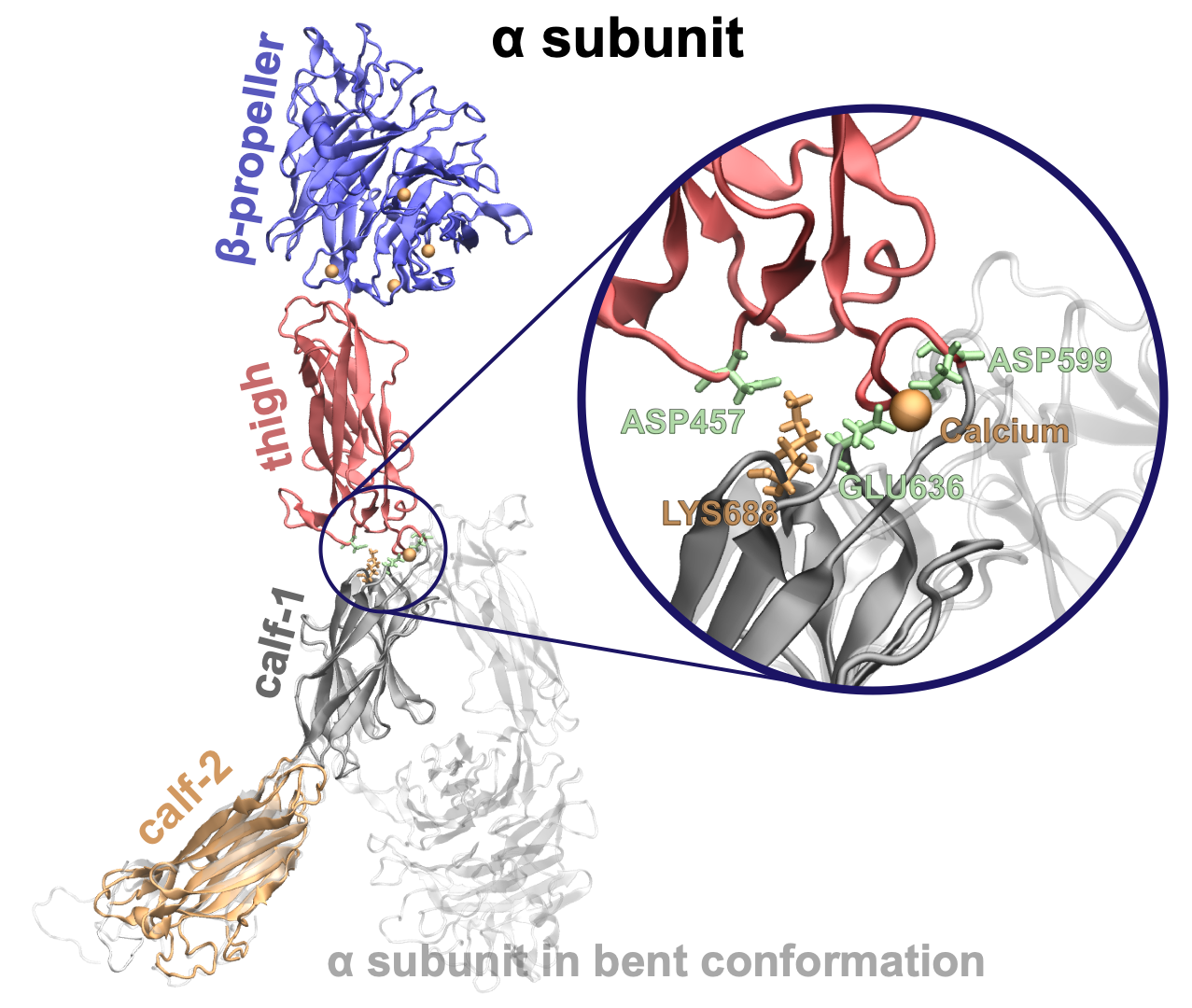

Supplementary Fig. 3. The salt bridge at the hinge in the α subunit.

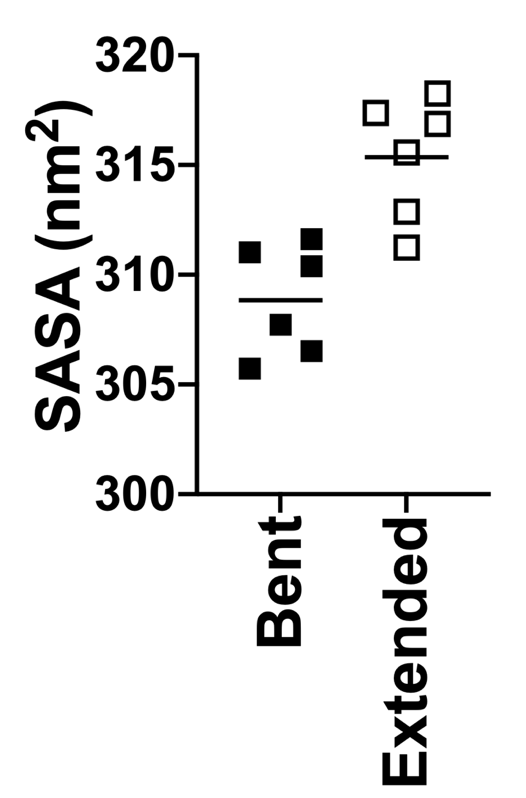

Supplementary Fig. 4. The SASA of the integrin in last 1 ns of 6 free dynamics simulations and 6 pulling simulations.
